# PhyloJunction: a computational framework for simulating, developing, and teaching evolutionary models

**DOI:** 10.1101/2023.12.15.571907

**Authors:** Fábio K. Mendes, Michael J. Landis

## Abstract

We introduce PhyloJunction, a computational framework designed to facilitate the prototyping, testing, and characterization of evolutionary models. PhyloJunction is distributed as an open-source Python library that can be used to implement a variety of models, through its flexible graphical modeling architecture and dedicated model specification language. Model design and use are exposed to users via command-line and graphical interfaces, which integrate the steps of simulating, summarizing, and visualizing data. This paper describes the features of PhyloJunction – which include, but are not limited to, a general implementation of a popular family of phylogenetic diversification models – and, moving forward, how it may be expanded to not only include new models, but to also become a platform for conducting and teaching statistical learning.

---

Phylogenetic models of lineage diversification have been applied to a wide variety of evolutionary phenomena spanning the disciplines of paleobiology [9, 26, 66], historical biogeography [8, 22, 39, 58], macroecology [11, 81], epidemiology [13, 51, 64], cancer evolution [41], molecular evolution [20, 76], and linguistics [27]. The evolutionary processes underlying these phenomena take place across a range of scales – from days to millions of years and from individual cells to the entire planet – and are known or hypothesized to operate under a similarly broad scope of tempos, modes, and spatial coordinates. Despite the heterogeneity in these biological phenomena, however, at the core of such phylogenetic models frequently lies the “state dependence” assumption: that the “states” of a lineage’s characters – ecological, geographic, phenotypic or genetic – may shape anagenetic and cladogenetic evolution. Stochastic processes that make such assumption, the so-called state-dependent speciation and extinction (SSE) processes, comprise a popular family of models [44] for the evolution of phylogenetic patterns.

In the recent past, many excellent methods for simulating under pure diversification models [e.g. 4, 24, 31, 43, 69] and SSE processes [e.g. 6, 19, 20, 43, 50] have been published. While overlapping in their capabilities, each of those methods was developed and uniquely optimized given a specific intended application; hence, they differ in terms of their model assumptions, implementation details and documentation, and execution attributes (e.g., speed, ease-of-use, etc.). Amidst this variation, we are unaware of any methods that, within a single cohesive codebase, can simultaneously (i) simulate under arbitrarily complex SSE scenarios (but see [78]), (ii) support an intuitive model specification grammar (e.g., [14, 30]), (iii) be easily extended by others to include new models, and (iv) showcase a built-in graphical user interface for automatic visualization and summarization of synthetic data, streamlining user interaction with the software (but see [14]).

In the hope of filling this gap in the computational biology toolbox, we introduce a new, open-source computational framework for evolutionary modeling: PhyloJunction. PhyloJunction ships with a very general SSE model simulator and with additional functionalities for model validation and Bayesian analysis. Importantly, we designed PhyloJunction around a graphical modeling architecture, and equipped it with a dedicated probabilistic programming language. These features are forward-looking; they will make it easy to expand and integrate PhyloJunction’s evolutionary model ecosystem in the future. PhyloJunction comes with a graphical user interface (GUI) that allows users to readily inspect and interact with simulation outputs, making this program amenable to classroom use. A command-line interface (CLI) is also available for running PhyloJunction remotely and in parallel.

## 1 Flexible simulation: prototyping, testing and characterizing evolutionary models

PhyloJunction was created first and foremost as an evolutionary model simulator, more specifically a flexible simulator of SSE diversification models. A series of related diversification models (Table 1) have been implemented in multiple computational methods with varying foci and performance, each making different assumptions about how a process starts and ends, whether it flows backward or forward in time, and what output is processed and presented to the user. PhyloJunction was born out of the necessity of coalescing the strengths of these different implementations in a single, cohesive application with additional capabilities (see below). As illustrated in later sections and in the online documentation, our implementation can simulate arbitrarily complex SSE processes (all models in Table 1; validation against other software can be found in the supplement), and presents the user with a variety of textual and graphical outputs.

**Table 1:** Phylogenetic models that can be simulated with PhyloJunction. Some of these models are nested within each other (e.g., BiSSE is a special case of MuSSE). “skyline” indicates time-heterogeneous rates varying in a piecewise-constant manner. “fossilized” means the addition of a fossilization parameter, which allows for direct ancestors in the reconstructed (sampled) tree. All models can be simulated with incomplete sampling. Representative papers for each model are listed under references.

| Model | A.k.a. or acronym | Reference(s) |
| --- | --- | --- |
| Pure-birth | Yule | [67, 83] |
| skyline |  |  |
| fossilized |  |  |
| Birth-death |  | [36, 52] |
| skyline | BDSKY | [72] |
| fossilized | FBD | [26] |
| Binary SSE | BiSSE | [44] |
| skyline |  |  |
| fossilized |  |  |
| Multistate SSE | MuSSE | [19] |
| skyline |  |  |
| fossilized |  |  |
| Geographic SSE | GeoSSE | [17] |
| skyline |  |  |
| fossilized |  |  |
| Cladogenetic SSE | ClaSSE | [22] |
| skyline |  |  |
| fossilized |  |  |
| Multitype birth-death | MTBD | [71] |
| skyline |  | [64] |
| fossilized |  |  |

Beyond its immediate goal of simulating SSE processes, however, it was evident early on that PhyloJunction could grow and serve more broadly as a computational framework for developing evolutionary models. This ultimate purpose manifests from PhyloJunction’s graphical model architecture being written in Python – a design and language convenient for prototyping model code, on which we expand below – and from the critical role simulation plays in model testing and characterization, two key stages in a model’s life cycle. A newly implemented model prototype is typically pitched against data simulated in simple scenarios, with the expectation that it returns acceptable parameter estimates given some truth (i.e., a value used in simulation). Upon failure, development loops back to implementation so bugs can be patched; this potentially iterative process is the testing (or validation) stage. If testing succeeds, the model is released to the public and enters a final characterization stage, in which its behavior and adequacy are thoroughly scrutinized by the scientific community, again via analysis of simulated data (e.g., [12, 40, 45, 46, 59, 63, 68]).

In the following sections, we detail the different features that allow PhyloJunction to flexibly specify and simulate diversification processes, and to facilitate the different steps involved in computational evolutionary model development.

## 2 A graphical model architecture and dedicated language for specifying arbitrarily complex models

Any type of analytical or generative procedure involving statistical models requires some form of infrastructure for specifying such models. One example is the framework adopted by the BEAST, BEAST 2 and RevBayes platforms, whereby atomic model components can be combined into an arbitrarily large Bayesian network – a probabilistic graphical model whose structure can be represented by a directed acyclic graph (DAG; or more explicitly as a factor graph, e.g., Fig. 1b; [29]). The popularity of these platforms is elevating graphical models to a modeling standard, although every one of these programs differs in how it allows users to specify models.

**Figure 1.**
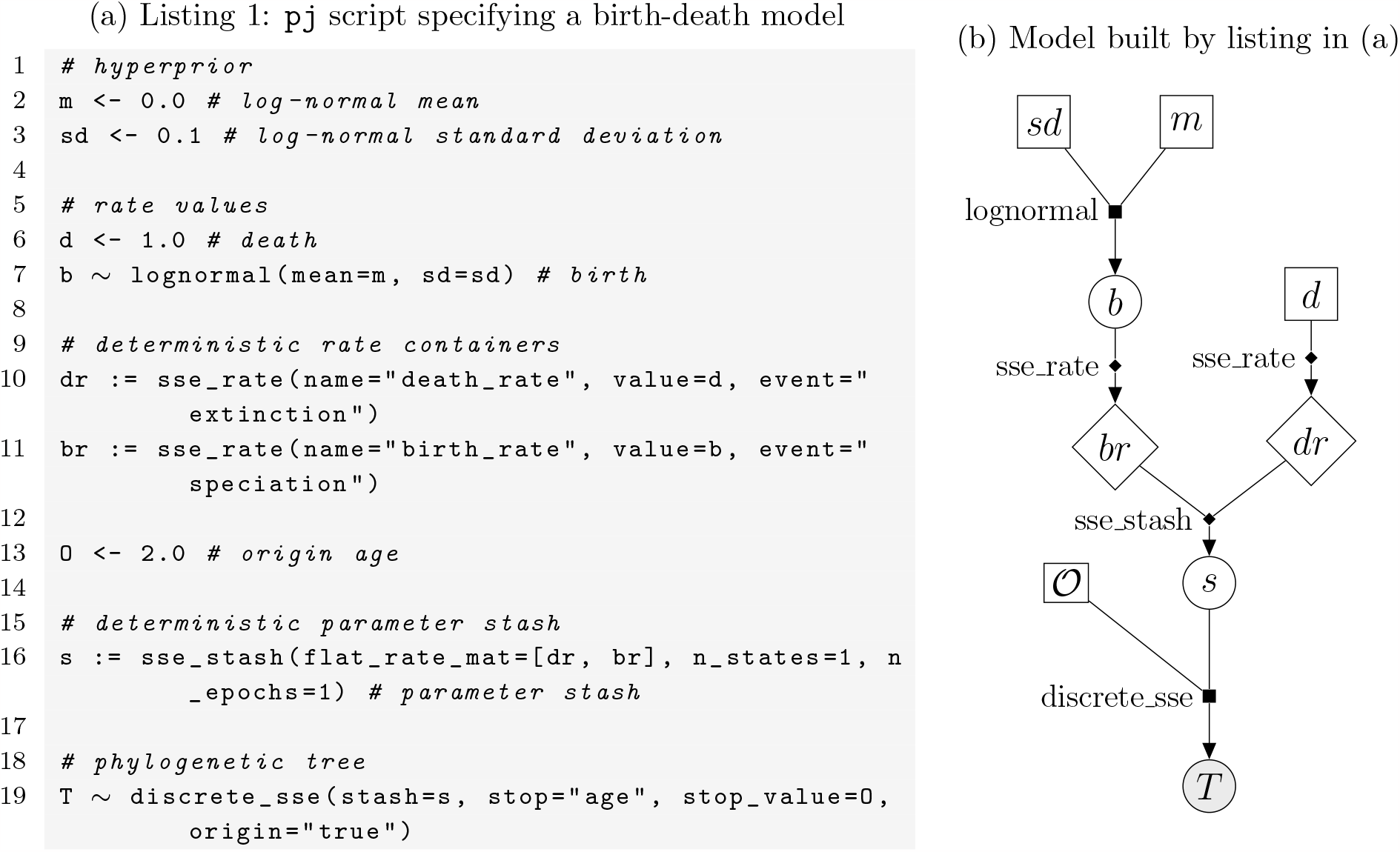
A birth-death phylogenetic model (a) as specified with phylojunction, PhyloJunction’s eponymous programming language, and (b) shown as a factor graph, a generalization of a directed acyclic graph (DAG). A few of the symbols in (b) were introduced in the context of phylogenetics by [29]. Briefly, empty squares and empty circles drawn in continuous lines represent constant and stochastic nodes, respectively. Circles filled in gray represent stochastic nodes whose values are observed (i.e., data). Empty diamonds denote deterministic nodes (the output of deterministic functions). Factors capture the conditional dependencies between stochastic nodes and are either (i) filled squares, each associated to a distribution characterized by a density function and from which values can be sampled, or (ii) filled diamonds, denoting each a deterministic function.

Here, we take a model specification approach that sits between those adopted by the BEAST and RevBayes community. PhyloJunction implements a programming language, `phylojunction` (written in lowercase and abbreviated as `pj`), together with an interactive development environment for specifying phylogenetic models (see the next section). `pj` is lightweight like popular markup languages (e.g., XML, BEAST’s format of choice), but resembles model scripting languages (e.g., Rev, the language introduced by RevBayes) in its syntax, hence its retained human-readability.

Like the Rev language, `pj` commands can be read as mathematical statements, and are naturally interpreted as instructions for building a node in a DAG (see below). User commands instruct the application engine to take some form of user input, produce some value from it, and then store that value permanently in a new variable created on the spot. Every command string consists of an assignment operator placed between the variable being created (on its left side) and some user input (on its right side). Listing 1 (Fig. 1a) demonstrates the different ways in which this essential operation takes place as a time-homogeneous birth-death model is specified.

Following the grammar of Rev [30], the behavior of a variable is determined by which assignment operator (<-, ∼, or :=) is used for assignment. For example, line 6 in listing 1 (Fig. 1a) creates a variable ‘d’ (the death rate), which is then passed and henceforth stores an unmodified user input, constant value 1.0. This type of constant value assignment is carried out with the constant assignment operator, ‘<-’. Line 7, in turn, shows how the stochastic assignment operator ‘∼’ is used to create a variable named ‘b’ (the birth rate). This variable will then store a random value drawn from a user-specified distribution. Here, the user input consists of the moments of a log-normal distribution.

Finally, the deterministic assignment operator, ‘:=‘, is used to assign a value computed deterministically from existing variables (or other user input) to a new variable. This is illustrated by lines 10, 11 and 16 in listing 1 (Fig. 1a). The purpose of deterministic assignments is to transform, combine or annotate one or more existing variables, and give users more control over model building. Without this class of explicit operations, such steps would instead take place out of sight in the backend, or alongside many other actions upon a single `pj` command, both of which can contribute to obscuring model structure.

Computer variables created with `pj` are nodes in the DAG that describes all variable dependencies, distributions, functions, and values that comprise the full evolutionary model. With every `pj` command the DAG thus grows by a node, which is immediately assigned a value. The nature of the assignment (constant, stochastic, or deterministic) reflects which operator was used, as explained above. A thorough treatment of the grammar and usage of graphical models for evolutionary inference can be found in [29, 30] and the tutorials therein.

### Technical remarks on the phylojunction language

In PhyloJunction, models are specified through commands written in the eponymous custom language, `phylojunction` (`pj`). In the current version of `pj`, created variables are the sole, immutable output of every function – and this output depends exclusively on a function’s arguments. Variable immutability has two consequences. First, it precludes loop control structures (e.g., *for* and *while* loops), with replication and “plating” (see [29]) being achieved instead through vectorization, a concept R users should be familiar with. (`pj` also does not support structures such as *if-then-else* and *switch* statements, effectively abstracting control flow.) Second, apart from the logical dependencies between nested functions – which reflect dependencies among DAG nodes – command evaluation order does not affect model specification and simulation. For example, in listing 1 (Fig. 1a), commands on lines 2, 3 and 6 are order-interchangeable, and so are those on lines 10 and 11, but the command on line 7 must be executed before that on line 11.

The features described above make `pj` behave largely as a declarative language like XML. While commands in `pj` are Rev-like in syntax, and instantiate and store a DAG object in memory (the state of a PhyloJunction section), the similarities with Rev end here. In contrast to an imperative scripting language (e.g., R, Python, Rev), `pj` (i) is easier to learn, understand and write, (ii) enhances reproducibility, (iii) leaves less room for programming mistakes (e.g., variable overwriting, container indexing errors), and (iv) shifts the user’s attention from how to specify a model to the structure of the model itself. Focus on model structure in PhyloJunction is further encouraged by `pj`’s grammar ignoring actions and settings unrelated to model building, such as dependency loading, input/output, Bayesian proposals, MCMC parameters, etc. All of these properties make `pj` a lightweight language that can be particularly useful in the classroom.

## 3 Standalone command-line and graphical user interfaces

PhyloJunction integrates its multiple utilities for simulating, testing and characterizing evolutionary models via both command-line (CLI) and graphical user interfaces (GUI; Fig. 2a). Through the CLI and GUI, users can provide PhyloJunction with a series of DAG-building instructions in the form of a `pj` script (e.g., listing 1, Fig. 1a). Users can also build a DAG by entering commands through the GUI’s command prompt (Fig. 2a, number 1). Synthetic data is then generated while a `pj` script (or sequence of commands) is processed, and can later be exported as text files to a user-specified location. The interfaces can be further used to save and load a particular model instance as a serialized byte stream.

**Figure 2:**
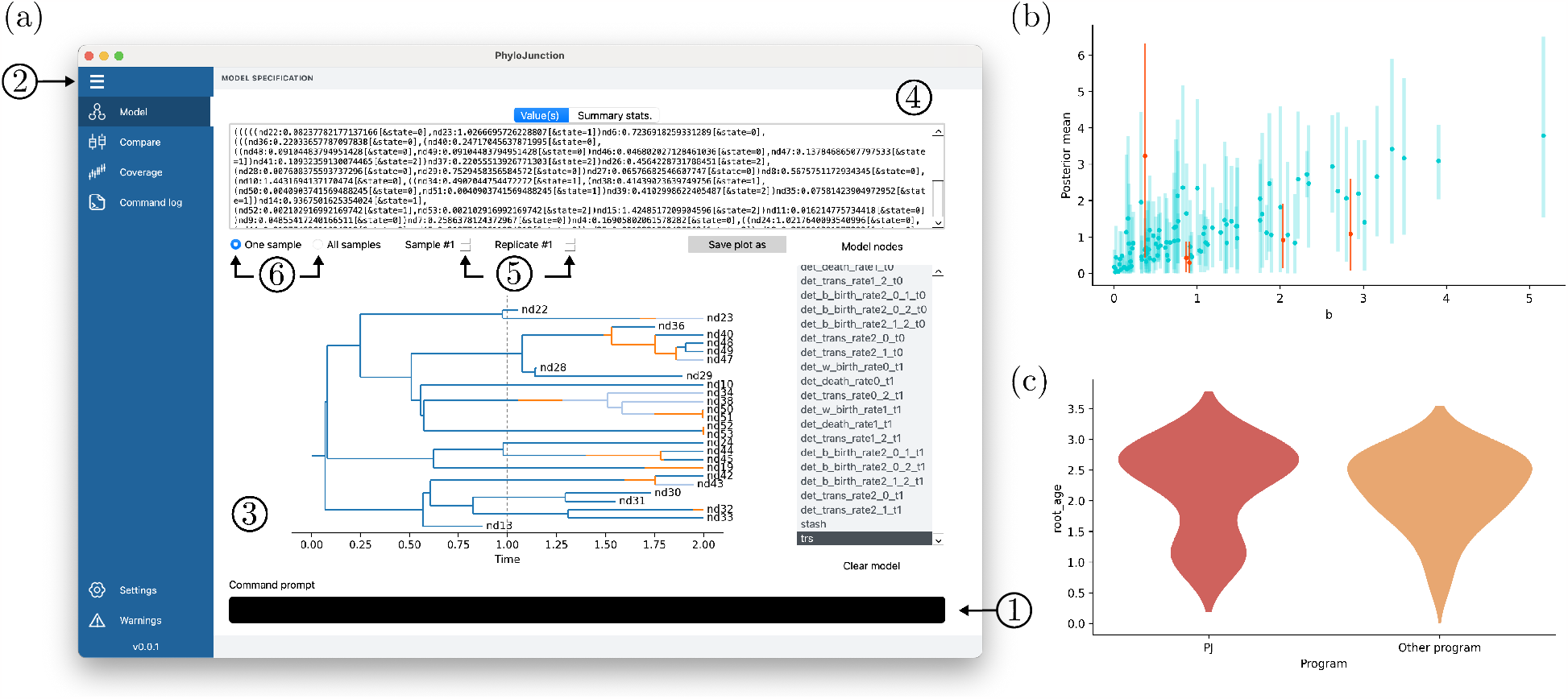
PhyloJunction’s (a) graphical user interface (GUI) with different features indicated by numbers (see main text), and plots from (b) “Coverage” and (c) “Compare” graphical exploration functionalities.

As any modern computer application, PhyloJunction’s GUI exposes its features to users via a menu (Fig. 2a, number 2). On the main tab (“Model”), one can navigate the DAG and see its node values as a plot, text string, or both (Fig. 2a, numbers 3 and 4). Users can also cycle through replicated simulations (Fig. 2a, number 5), and examine node-value summaries computed for individual simulations or across replicates (Fig. 2a, number 6). Node-value summaries include the mean and standard deviation for scalar variables, and statistics like the root age and number of tips for phylogenetic trees.

Automatic summarization and visual inspection of synthetic data expedites model testing and characterization, by helping researchers quickly determine if a model setup is appropriate. Empiricists can promptly examine the effect that prior choice may have during Bayesian inference, for example, depending on what simulated data sets look like. Under an SSE model [13], high state-transition rates causing saturation would be immediately discernible in the state mappings coloring a phylogenetic tree (Fig. 2a, number 3).

### Workflow functionalities for model testing, characterization, and teaching

In addition to data simulation, the different stages of model development have a few common denominators. Researchers must usually contend with (i) parsing inference results, (ii) comparing parameter estimates and their true values, and (iii) displaying the results as graphs. These tasks will commonly be repeated across parameter space, and sometimes under different models altogether. Furthermore, testing and characterization pipelines are often built, executed and described several times by multiple researchers – even when the procedures taking place are very similar (e.g., [28, 48]). This redundancy is not only an inefficient use of researchers’ time, but also hinders reproducibility.

PhyloJunction introduces a suite of utilities for streamlining and automating model specification, testing and characterization. These are meant to minimize scripting redundancy and maximize the reproducibility of *in silico* experiments. Different utilities are separately documented and can be invoked by the user from within custom Python scripts, as modules. Alternatively, users may access PhyloJunction’s features via its standalone interfaces.

Validation utilities, for example, can be accessed via the GUI’s “Coverage” tab. These were designed with Bayesian coverage validation in mind (e.g., [21, 53, 84]). When a simulated data set is analyzed using a Bayesian platform, this tab can be used for loading raw or post-processed inference result files. True parameter values must be loaded as a table, or if data was simulated with PhyloJunction, users can load a model instance (saved previously as a byte stream). Parameter coverage can then automatically computed, and Bayesian intervals plotted against true parameter values (Fig. 2b).

The GUI’s “Compare” tab, in turn, exposes additional model exploration utilities to the user. Here, parameter values generated by PhyloJunction under a model can be visualized against theoretical expectations, or against values simulated by a different program. Because PhyloJunction automatically computes parameter summary statistics, those can also be displayed side-by-side with comparable quantities calculated elsewhere (Fig. 2c). These functionalities are useful in a Bayesian context, for example, whenever a model has been implemented for inference, but not for direct simulation. In such cases, one can use PhyloJunction to rapidly build a direct simulator for the first time, and then use the “Compare” tab to check it against Monte Carlo samples produced by the existing implementation.

Simulation functionalities as well as those available under the “Coverage” and “Compare” tabs were developed because of the ubiquitous (and repetitious) nature of certain tasks involved in validating and characterizing a model. In addition to methodological research, however, we anticipate that these features will find use in teaching settings – especially considering the growing availability and popularity of technical workshops [2, 3], and new pedagogical material [25, 62, 80]. While trying their hand at implementing simple evolutionary models, students could use PhyloJunction to validate said models or to obtain simulation benchmarks, and to immediately visualize results via the GUI. PhyloJunction’s pedagogical impact will be further enhanced by its implementation in pure Python (see below), a cross-platform, user-friendly language that finds widespread use in the classroom.

## 4 Longevity through an extensible and user-friendly model ecosystem

One hurdle that must be often overcome during model development is the steep learning curve of the low-level programming languages many software platforms are written in. RevBayes is written in C++, for example, while BEAST and BEAST 2 are developed in Java. This is a choice motivated by compiled programming languages generally outperforming interpreted languages (e.g., R, Python), and being preferred over the latter whenever speed is a priority, such as when a method is primarily used for inference from challenging data sets. Languages like C++ and Java also natively support object-oriented programming – a programming paradigm that is critical for erecting vast, extensible and maintainable codebases such as those living inside those platforms.

Despite being conversant in interpreted languages, many biologists with an enthusiasm for evolutionary modeling have little to no experience with the commonly abstruse syntax and features of low-level languages (e.g., memory management, abstraction, typing). They also have rarely had to contend with the complicated pipelines for compiling large programs across different types of computers, and with the configuration of industry-grade IDEs (integrated development environment), used for navigating immense codebases. Unless working closely with developers of big software platforms, individual scientists are likely to struggle with (i) reverse-engineering complex code that may not have been written to be read by others, and (ii) adding new code that does not break the behavior of the original codebase.

One alternative that obviates some of these difficulties is to implement and release models as R or Python packages (e.g., [4, 10, 19, 21, 50, 56, 61]). A package has a comparatively small codebase that can be written by anyone from scratch, is self-contained and thus easily maintainable, and can be integrated with other packages more or less readily, via the scripting language. Furthermore, public package archives such as CRAN (the Comprehensive R Archive Network) or PyPI (the Python Package Index) do not restrict how a package should be programmed; package source files are immensely variable in their coding language, conventions and documentation, and programming paradigm. The minimal package submission and code-design requirements of CRAN and PyPI allow researchers freedom and flexibility, both unquestionable advantages to this variety of method development.

Writing packages has its challenges. Developers who want to add to or combine existing packages will likely have to contend with code written in a mix of languages (e.g., R, Python, C, C++, FORTRAN), paradigms (e.g., functional, objected-oriented) and styles. Furthermore, CRAN and PyPI put the onus on the researcher to choose among (often multiple) packages for the same or different purposes. Packages may vary with respect to their underlying algorithms, modeling assumptions and notation (see [70] for an example). Lastly, every scientist will adopt a unique R scripting strategy when specifying a model. All of the above makes reproducibility of results harder, and leads to code that is often chimeric (in its style, paradigm and language), single-use, or redundant.

The choice of platform for writing modeling software thus involves trade-offs related to technical difficulty, speed, distributability, and maintainability. PhyloJunction embodies our attempt at balancing the above considerations while introducing an alternative methodology for model development and characterization. The brunt of PhyloJunction’s design effort involved conceiving a computational framework that could not merely be extended – among other things, our intention is to facilitate the early stages of model prototyping and testing – but extended with minimal refactoring and in the most developer-friendly way possible.

We chose to implement Phylojunction in Python primarily because of its native support for object-oriented programming, a paradigm that aids codebase expansion and maintenance. Furthermore, Python has clear community standards and many tools (e.g., *mypy, Sphinx, pep8* [77], *pep20* [57]) for encouraging or enforcing conventions on coding style, type hinting and documentation – all of which further contribute to codebase clarity and consistency. A Python codebase can also be easily navigated with any of the various user-friendly IDEs with support for Python (e.g., Visual Studio Code, PyCharm, Spyder).

Finally, Python development can profit from a vast array of free, industry-grade scientific libraries for data manipulation (e.g., *matplotlib* [34], *pandas*), statistics and Bayesian analysis (e.g., *scipy* [79], *PyMC3* [35], *ArviZ* [38]), and machine learning (e.g., *TensorFlow* [1], *scikit-learn* [55]). Of particular relevance to PhyloJunction’s is PyPI’s growing list of modules specifically aimed at phylogenetic or population genetic analysis (e.g., *DendroPy* [73], *PyRate* [65], *MESS* [54], *ete3* [33], *msprime* [5]), some of which have already been or may be integrated with PhyloJunction in the future. Below we suggest a few ways in which the latter may be done.

## 5 Availability and resources

PhyloJunction’s source code is publicly available on https://github.com/fkmendes/PhyloJunction. Documentation on how to install and use the program can be found on https://phylojunction.org. PhyloJunction is licensed under GNU General Public License v3.0.

## 6 Future directions

We introduced PhyloJunction, an open-source package for simulating state-dependent speciation and extinction (SSE) processes, a large family of diversification models that has found success across a range of scientific domains [13, 27, 75]. Most implementations of SSE models have prioritized inference and efficiency over simulation and generality; the latter is the relatively vacant niche PhyloJunction was designed to fill. In addition to model-specification and simulation tools, our program ships with a series of utilities for summarizing and visualizing simulation outputs, as well as data-wrangling functions for model validation and characterization. These utilities are integrated and exposed to users by standalone command-line and graphical interfaces, which simplify the execution and reproduction of in silico experiments.

Models in PhyloJunction are embedded within a graphical modeling architecture, which also underlies the package’s dedicated probabilistic-programming language, phylojunction. These features make PhyloJunction’s model ecosystem extensible beyond SSE processes, and allow its components to be promptly integrated. Future software releases are planned to include distributions for different types of data models (e.g., DNA and protein sequences, [23, 74, 82]; discrete and continuous characters, [18, 42]), evolutionary clock models (e.g., [15, 16]), population-genetic and phylogeographic processes (e.g., [37, 60]), or models combining any of the above (e.g., [7, 32, 47]). A richer selection of evolutionary processes should widen the range of potential applications of PhyloJunction in research and teaching.

PhyloJunction was primarily designed to be a framework for simulation and prototyping of evolutionary models, but we expect its future development to further take on the task of statistical inference. Moving in that direction may involve introducing subroutines for creating textual instructions for Bayesian inference, for example, as required by popular platforms (e.g., RevBayes, BEAST, BEAST 2). Bayesian inference is also possible within a Python environment, although it is unclear how immediately useful existing libraries (e.g., [35]) may be in terms of parameter estimation in phylogenetic space. Porting or implementing Bayesian inference utilities in our platform would in the very least allow synthetic data to be simulated under simple models, and immediately plotted. Such an extension would further empower PhyloJunction as a teaching tool. Alternatively, it should be straightforward to integrate PhyloJunction’s functionalities for summarizing data together with Python machine learning libraries. There is increasing evidence [49] backing machinelearning methods as viable alternatives to frequentist and Bayesian evolutionary inference, especially when the latter is very onerous or impossible [75].

It is our long-term hope for PhyloJunction that it not only increasingly facilitates research in evolutionary modeling, but that its capabilities can be diversified and enhanced by (and according to the needs of) the scientific community at large.

## 7 Acknowledgements

We thank Felipe Zapata, Isaac Lichter-Marck, Albert Soewongsono, Sarah Swiston, Sean McHugh, and Ammon Thompson for conversations that motivated the development of PhyloJunction, inspired its design, and improved its manuscript.

