## Supplementary material for "PhyloJunction: a computational framework for simulating, developing, and teaching evolutionary models"

Fábio K. Mendes[1,*] and Michael J. Landis[1]

[1] *Department of Biology, Washington University in St. Louis, St. Louis, MO*

### 1 Comparisons with other software

`PhyloJunction`'s (PJ) simulation code was compared to independently implemented counterparts whenever possible, which in some cases included multiple software packages. The latter included packages written in R, such as `geiger` [3], `diversitree` [2], `phytools` [4], `TreeSim` [5], and `FossilSim` [1], as well as in Java, such as `MASTER` [6]. As mentioned in the main text, each of those tools is unique in its conditioning of diversification models and filtering of simulated output.

Comparisons under different models (Supplementary Figs. 1, 2, 3, 4, 5, and 6) were carried out in multiple arbitrary regions of parameter space (Supplementary Tables 1 to 4). Deciding which programs to compare in each scenario was largely determined by our perceived ability to match the model assumptions and output parsing of different simulators.

[1] PJ implements the "simple sampling approach" (SSA; see Stadler, 2011)

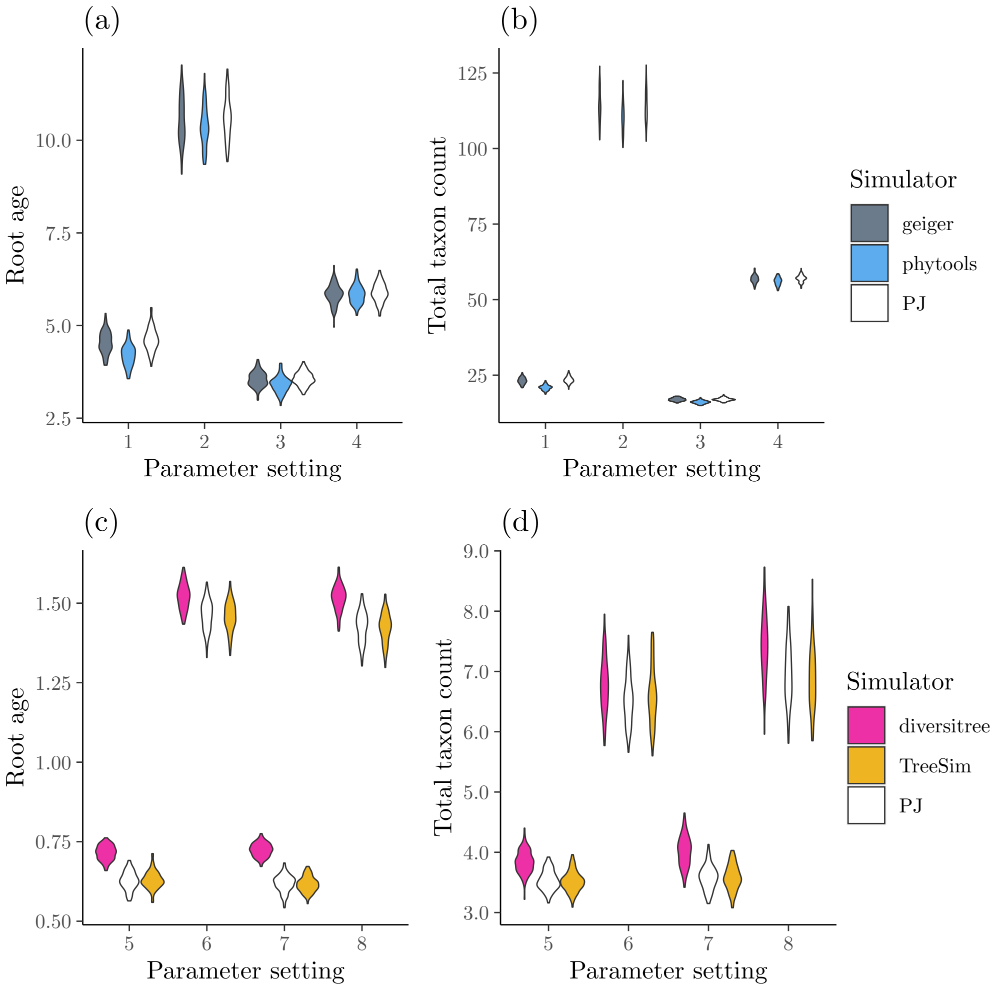


Supplementary Figure 1: Summaries of data sets simulated under the birth-death model with `PhyloJunction` (PJ), and the `geiger`, `phytools` and `diversitree` R packages. Quantities are summarized from complete (i.e., including extinct taxa) trees, assuming perfect sampling. Summaries include (a, c) root ages, and (b, d) total taxon count. Each violin plot comprises 100 values, each corresponding to the focal statistic (e.g, root age) averaged over 100 trees. Parameter settings are detailed in Supplementary Table 1.

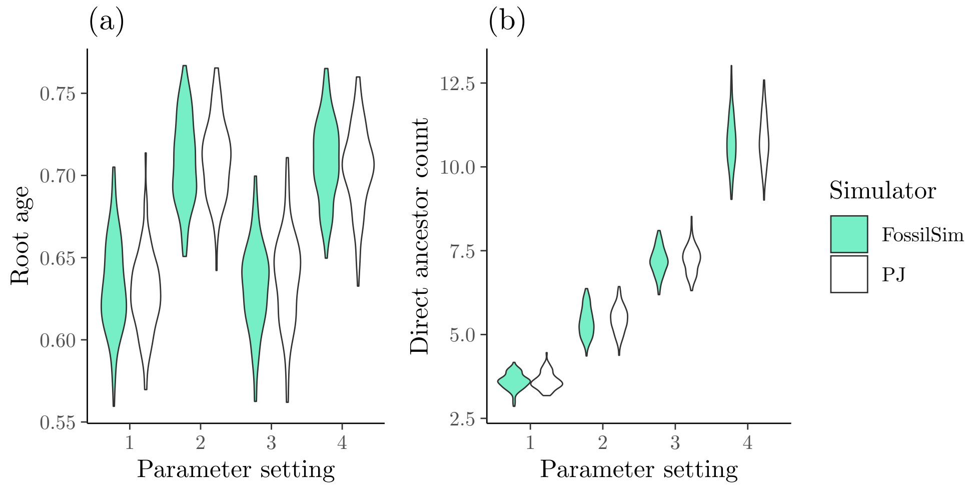


Supplementary Figure 2: Summaries of data sets simulated under the fossilized birth-death (FBD) model with `PhyloJunction` (PJ) and the `FossilSim` R package. Quantities are summarized from complete (i.e., including extinct taxa) trees, assuming perfect sampling. Summaries include (a) root ages, and (b) direct ancestor taxon count. Each violin plot comprises 100 values, each corresponding to the focal statistic (e.g, root age) averaged over 100 trees. Parameter settings are detailed in Supplementary Table 1.

Supplementary Table 1: Model configurations used in PhyloJunction validation. Dots denote parameters that do not apply to a given model. "BD" stands for birth-death, "FBD" for fossilized birth-death and "BiSSE" for binary state-dependent speciation and extinction models.

| Tree model | Parameter setting | $\lambda$ or $\lambda_0, \lambda_1$ | $\mu$ | $\psi$ | $q_{01}$, $q_{10}$ | Max. age | Max. extant taxon count[1] | Start from |
|---|---|---|---|---|---|---|---|---|
| BD | 1 | 1.0 | 0.8 | · | · | · | 10 | Root |
| | 2 | 1.0 | 0.8 | · | · | · | 30 | Root |
| | 3 | 1.0 | 0.5 | · | · | · | 10 | Root |
| | 4 | 1.0 | 0.5 | · | · | · | 30 | Root |
| | 5 | 1.0 | 0.8 | · | · | 1.0 | · | Origin |
| | 6 | 1.0 | 0.8 | · | · | 2.0 | · | Origin |
| | 7 | 1.0 | 0.5 | · | · | 1.0 | · | Origin |
| | 8 | 1.0 | 0.5 | · | · | 2.0 | · | Origin |
| FBD | 1 | 1.0 | 1.0 | 2.0 | · | 1.0 | · | Origin |
| | 2 | 2.0 | 1.0 | 2.0 | · | 1.0 | · | Origin |
| | 3 | 1.0 | 1.0 | 4.0 | · | 1.0 | · | Origin |
| | 4 | 2.0 | 1.0 | 4.0 | · | 1.0 | · | Origin |
| BiSSE | 1 | 1.0, 0.75 | 0.5 | · | 0.25, 0.75 | 5.0 | · | Origin |
| | 2 | 1.0, 0.75 | 0.5 | · | 0.5, 0.5 | 5.0 | · | Origin |
| | 3 | 1.0, 0.75 | 0.5 | · | 0.75, 0.25 | 5.0 | · | Origin |
| | 4 | 1.0, 0.75 | 0.5 | · | 1.0, 0.0 | 5.0 | · | Origin |

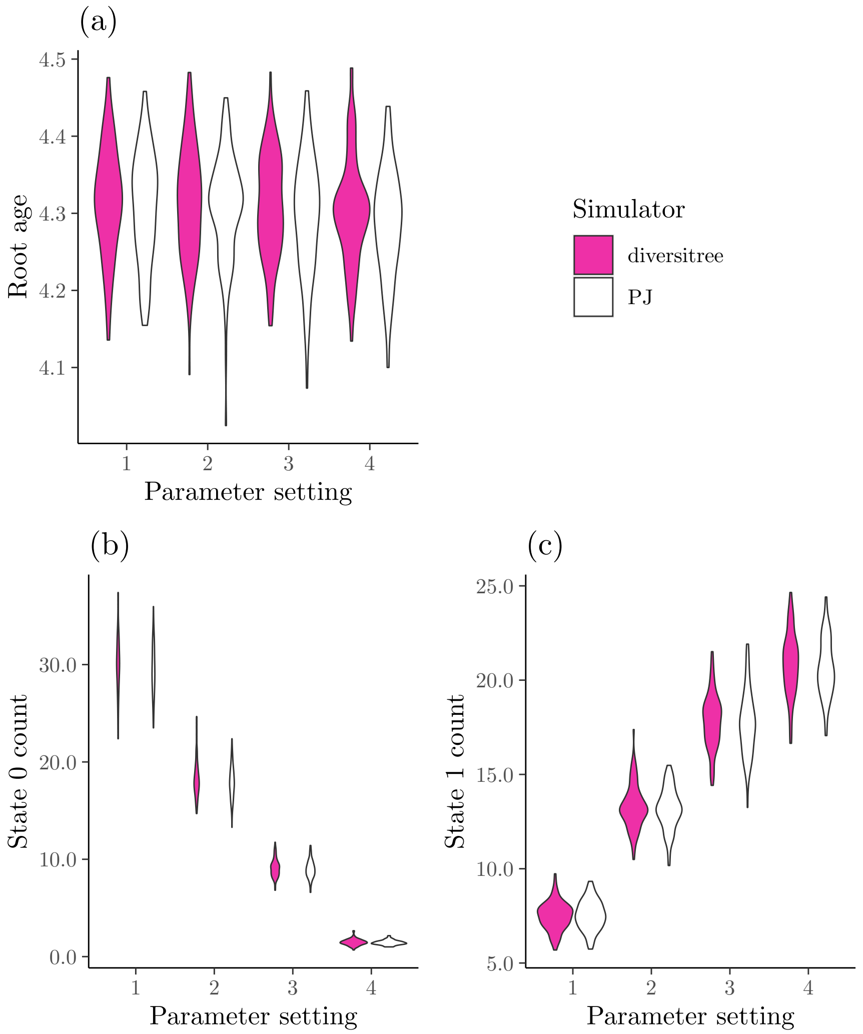


Supplementary Figure 3: Summaries of data sets simulated under the binary state-dependent speciation and extinction (BiSSE) model with `PhyloJunction` (PJ) and the `diversitree` R package. Quantities are summarized from complete (i.e., including extinct taxa) trees, assuming perfect sampling. Summaries include (a) number of taxa at state 0, (b) number of taxa at state 1, and (c) root ages. Each violin plot comprises 100 values, each corresponding to the focal statistic (e.g, root age) averaged over 100 trees. Parameter settings are detailed in Supplementary Table 1.

Supplementary Table 2: Geographic state-dependent speciation and extinction (GeoSSE) model configurations used in PhyloJunction validation. Processes started at the origin, stopped at a maximum age of age of 4.0, and were conditioned on the survival of at least one living taxon. (parameter names within parentheses follow 'diversitree"s notation).

| Tree model | Parameter setting | $\lambda_1$ (sA) | $\lambda_2$ (sB) | $\lambda_{0,1,2}$ (sAB) | $\mu_1$ (xA) | $\mu_2$ (xB) | $q_{1,0}$ (dA) | $q_{2,0}$ (dB) |
|---|---|---|---|---|---|---|---|---|
| GeoSSE | 1 | 1.25 | 1.25 | 0.75 | 1.0 | 1.0 | 1.0 | 1.0 |
| | 2 | 1.25 | 1.25 | 1.0 | 1.0 | 1.0 | 1.0 | 1.0 |
| | 3 | 1.25 | 1.25 | 1.25 | 1.0 | 1.0 | 1.0 | 1.0 |
| | 4 | 1.25 | 1.25 | 1.5 | 1.0 | 1.0 | 1.0 | 1.0 |

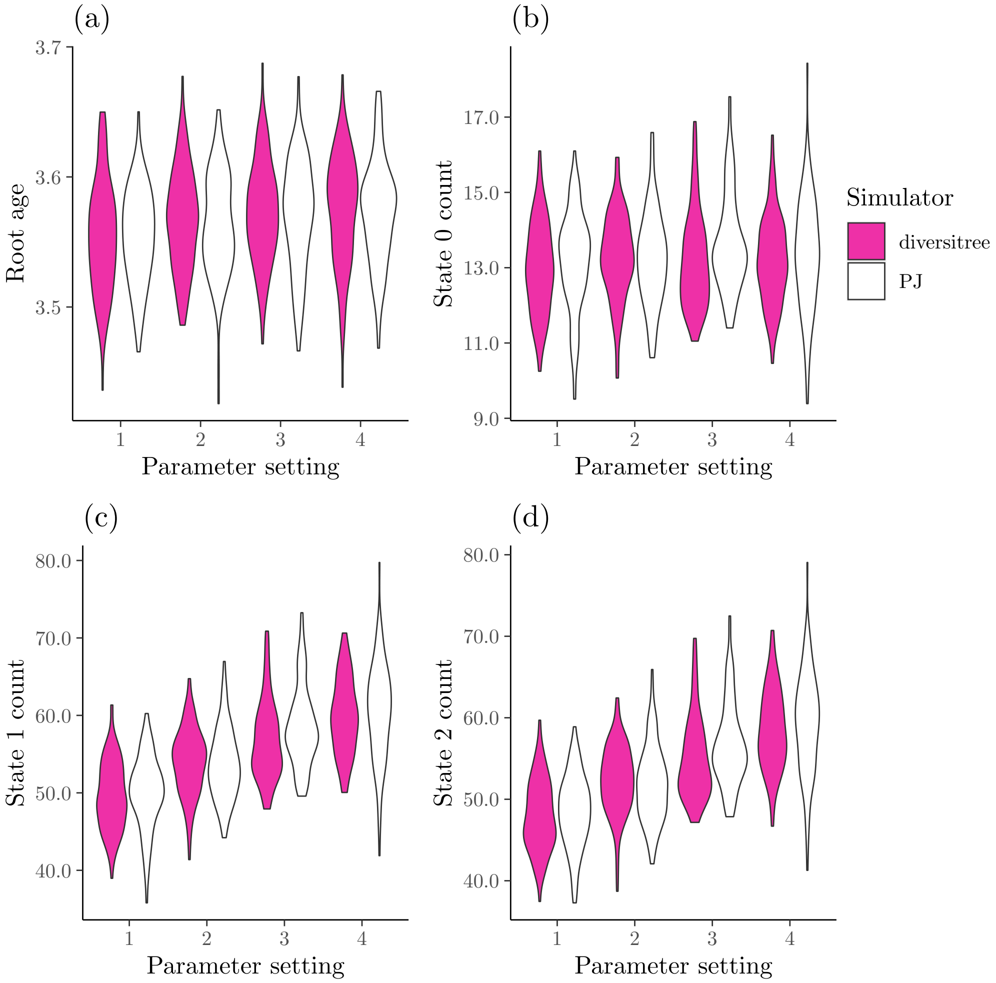


Supplementary Figure 4: Summaries of data sets simulated under the geographic state-dependent speciation and extinction (GeoSSE) model with `PhyloJunction` (PJ) and the `diversitree` R package. Quantities are summarized from complete (i.e., including extinct taxa) trees, assuming perfect sampling. Summaries include (a) root ages, (b) number of taxa at state 0, (c) number of taxa at state 1, and (d) number of taxa at state 2. Each violin plot comprises 100 values, each corresponding to the focal statistic (e.g, root age) averaged over 100 trees. Parameter settings are detailed in Supplementary Table 2.

Supplementary Table 3: Time-heterogeneous Yule model configurations used in PhyloJunction validation. All model configurations specified processes starting at the origin, and stopping at a maximum age of 6.0. All epoch starting times $t$ are defined in forward time units, with $t_0$, $t_1$ and $t_2$ corresponding to the starts of the first, second and third epochs, respectively. Each epoch, from oldest to youngest, was specified its own birth-rate, $\lambda^0$, $\lambda^1$, and $\lambda^2$.

| Tree model | Parameter setting | $t_0$ | $t_1$ | $t_2$ | $\lambda^0$ | $\lambda^1$ | $\lambda^2$ |
|---|---|---|---|---|---|---|---|
| Yule | 1 | 0.0 | 1.0 | 2.0 | 2.0 | 0.5 | 0.1 |
| | 2 | 0.0 | 1.0 | 3.0 | 2.0 | 0.5 | 0.1 |
| | 3 | 0.0 | 1.0 | 4.0 | 2.0 | 0.5 | 0.1 |
| | 4 | 0.0 | 1.0 | 5.0 | 2.0 | 0.5 | 0.1 |

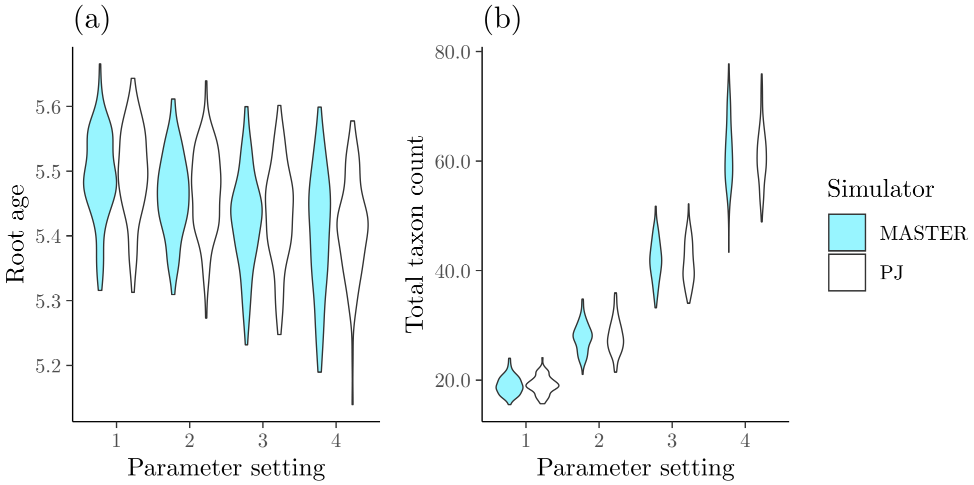


Supplementary Figure 5: Summaries of data sets simulated under a time-heterogeneous Yule model with `PhyloJunction` (PJ) and the `MASTER` BEAST 2 package. Quantities are summarized from perfectly sampled trees. Summaries include (a) root ages, and (b) total taxon count. Each violin plot comprises 100 values, each corresponding to the focal statistic (e.g, root age) averaged over 100 trees. Parameter settings are detailed in Supplementary Table 3.

Supplementary Table 4: Time-heterogeneous binary state-dependent speciation and extinction (BiSSE) model configurations used in PhyloJunction validation. All model configurations specified processes starting at the origin, and stopping at a maximum age of 6.0. All epoch starting times $t$ are defined in forward time units, with $t_0$ and $t_1$ corresponding to the starts of the first and second epochs, respectively. Birth-rates and transition rates were kept contant across epochs. Each epoch, from oldest to youngest, was specified its own death-rate for state 1, $\mu_1^1$.

| Tree model | Parameter setting | $t_0$ | $t_1$ | $\lambda_0^0$ | $\lambda_0^0$ | $\mu_0^0$ | $\mu_1^0$ | $\mu_0^1$ | $\mu_1^1$ | $q_{0,1}^0$ | $q_{0,1}^0$ | $q_{0,1}^1$ | $q_{0,1}^1$ |
|---|---|---|---|---|---|---|---|---|---|---|---|---|---|
| BiSSE | 1 | 0.0 | 2.0 | 1.1 | 1.2 | 1.0 | 0.0 | 1.0 | 0.85 | 0.0 | 0.0 | 1.0 | 0.5 |
| | 2 | 0.0 | 3.0 | 1.1 | 1.2 | 1.0 | 0.0 | 1.0 | 0.9 | 0.0 | 0.0 | 1.0 | 0.5 |
| | 3 | 0.0 | 4.0 | 1.1 | 1.2 | 1.0 | 0.0 | 1.0 | 0.95 | 0.0 | 0.0 | 1.0 | 0.5 |
| | 4 | 0.0 | 5.0 | 1.1 | 1.2 | 1.0 | 0.0 | 1.0 | 1.0 | 0.0 | 0.0 | 1.0 | 0.5 |

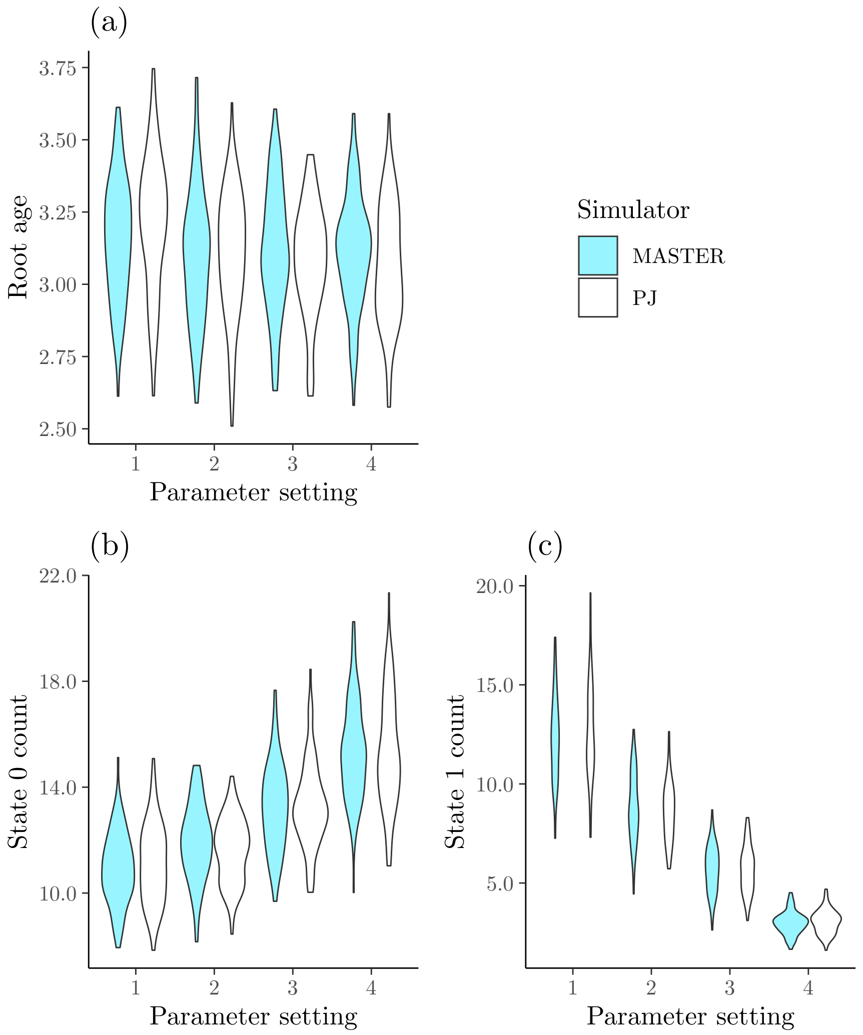


Supplementary Figure 6: Summaries of data sets simulated under a time-heterogeneous binary state-dependent speciation and extinction (BiSSE) model with `PhyloJunction` (PJ) and the `MASTER` BEAST 2 package. Quantities are summarized from complete, perfectly sampled trees. Summaries include (a) root ages, (b) number of taxa at state 0, (c) number of taxa at state 1. Each violin plot comprises 100 values, each corresponding to the focal statistic (e.g, root age) averaged over 100 trees. Parameter settings are detailed in Supplementary Table 4.
